# Reliability and correlates of intra-individual variability in the oculomotor system

**DOI:** 10.1101/535344

**Authors:** Marlou Nadine Perquin, Aline Bompas

## Abstract

Even if all external circumstances are kept equal, the oculomotor system shows intra-individual variability over time, affecting measures such as microsaccade rate, blink rate, pupil size, and gaze position. Recently, some of these measures have been associated with ADHD on a between-subject level. However, it remains unclear to what extent these measures constitute stable individual traits. In the current study, we investigate the intra-individual reliability of these oculomotor features. Combining results over three experiments (> 100 healthy participants), we find that most measures show good intra-individual reliability over different time points (repeatability) as well as over different conditions (generalisation). However, we find evidence against any correlation with self-assessed ADHD tendencies, mind wandering, and impulsivity. As such, the oculomotor system shows reliable intra-individual reliability, but its benefit for investigating self-assessed individual differences in healthy subjects remains unclear. With our results, we highlight the importance of reliability and statistical power when studying between-subject differences.

## Introduction

Imagine that you are working in your office, and one of your colleagues suddenly walks in: Your eyes will immediately change position from your work and will subsequently fixate on your colleague, and your pupil size will be modulated by the differences in light hitting your eye. These types of changes in eye position and pupil size may be described as ‘exogenous’ intra-individual variability – variability within an individual over time that is brought about by changes in the external environment. However, even when all external circumstances remain the same and one is solely fixating on a static dot, the eyes are still not ‘perfectly stable’. Instead, there will still be small changes in eye position (i.e., ‘fixational eye movements’, see Rolfs, 2009 for a review), small changes in pupil size, and blinks. All of these changes may be described as ‘endogenous’ intra-individual variability – brought about by internal fluctuations.

It seems reasonable that endogenous variability differs between individuals. Supporting this, a recent paper found positive associations between endogenous oculomotor variability and Attention-Deficit and/or Hyperactivity Disorder (ADHD) tendencies (Panagiotidi, Overton & Stafford, 2017). However, the intra-individual reliability of endogenous oculomotor variability is still largely unknown – meaning it is unclear to what extent this variability constitutes a reliable individual trait. While there is evidence for intra-individual reliability in oculomotor measures over different types of tasks (Andrews & Coppola, 1999; Boot, Becic & Kramer, 2009; Castelhano & Henderson, 2008; Poynter, Barber, Inman & Wiggins, 2013; Rayner et al., 2007), the properties of oculomotor variability during rest has not been much investigated. Such reliability is an important quality for any potential biomarker (Mayeux, 2004). The aim of the current paper is therefore twofold. First, we aim to examine whether variability in the oculomotor system shows reliable intra-individual consistency over different time points and different conditions in ‘resting state’ circumstances, to investigate to what extent this endogenous variability may be a reliable individual property. Secondly, we aim to investigate potential interindividual differences, by testing whether oculomotor variability correlates with ADHD, mind wandering, and impulsivity.

### Oculomotor functioning and variability

While ‘saccades’ refer to sudden, ballistic movements in eye position and ‘fixations’ refer to the maintenance of the eye position on a particular spot, microsaccades refer to small, sudden movements of the eye position during fixations (see Rolfs, 2009 for a review). Microsaccades are one of three types of fixational eye movements, the other two being drift and tremor. The movements of microsaccades have been described as ‘jerk-like’, small (typically being below 1-2° in amplitude), and often as ‘binocular’ (i.e., occurring in both eyes simultaneously). There have been several suggestions about the purpose of microsaccades (and fixational eye movements in general), relating to control over fixation position, prevention of perceptual fading, improvement of visual processing, (small-area) scanning of the environment, and acuity (see Rolfs, 2009; Martinez-Conde, Otero-Millan & Macknik, 2013 for reviews).

While microsaccades have been related to attention, this refers mostly to attentional cuing and ‘covert attention’ (i.e., foci of attention that are separate from the current eye position). Attentional cuing has been known to modulate both the direction and the occurrence of microsaccades, with the latter most commonly following a shape known as the ‘microsaccade rate signature’ – showing a sudden drop in microsaccades after cue onset, followed by a strong increase right after. Interestingly, this modulation of microsaccade rate seems influenceable by top-down expectations (Valsecchi, Betta & Turatto, 2007). However, the role of attentional cuing relates to exogenous variability, not to the manifestation of variability during rest – which would instead be related to fluctuations in internal states over time.

Intra-individual stability of oculomotor variability has been shown previously over different types of tasks, images, and display modalities (Andrews & Coppola, 1999; Boot et al., 2009; Castelhano & Henderson, 2008; Poynter et al., 2013; Rayner et al., 2007). For example, Castelhano & Henderson (2008) found consistency in individuals’ oculomotor behaviour between images in different display formats, but also between faces and scenes. In these cases, it seems plausible that the intra-individual consistency appears because of individual consistency in viewing and processing information; individuals may have a ‘default way’ of information processing that is reflected in their oculomotor behaviour. This is supported by findings that oculomotor behaviour can be altered when participants are given different instructions and feedback (Boot et al., 2009), and explains why it shows cultural differences (Rayner et al., 2007).

The intra-individual correlations of endogenous variability have furthermore been studied in relation to task-based variability. Andrews and Coppola (1999) looked at fixation duration and saccade size across five conditions: a ‘dark room’ condition, in which participants’ eye movements were continuously recorded for 100 seconds, two ‘viewing’ conditions, in which participants viewed simple and complex patterns, and two more ‘cognitive’ tasks, in which participants did visual search and reading. Oculomotor measures in the dark room condition showed positive intra-individual correlations to the viewing conditions, but not to cognitive conditions. Poynter et al. (2013) used a larger array of measures: For each participant, they extracted six measures of oculomotor activity (saccade amplitude, microsaccade rate and amplitude, and fixation rate, duration, and size) over four different tasks (a sustained fixation, scan-identify, search, and Stroop task). They found that each oculomotor measure correlated to itself between the different tasks within participants. However, their fixation task consisted of trials that were only three seconds long – meaning that the variability is still highly dependent on stimulus-onset, and that the task is not aimed at capturing (mostly) endogenous variability. Overall, none of these articles address the question of the current research directly – namely, to what extent endogenous variability itself is a reliable individual trait.

### Oculomotor variability and ADHD symptomatology

In the search for a potential ‘biomarker’ of ADHD, two previous studies investigated the relation between ADHD and oculomotor variability. Fried et al. (2014) examined differences between adults with ADHD (both in an ‘unmedicated’ and ‘medicated’ session) and healthy controls (unmedicated in both sessions). Participants were asked to make a button press in response to targets but not to non-targets. In their ‘unmedicated’ session, participants with ADHD showed significantly higher microsaccade and blink rates compared to controls, both near stimulus onset and throughout the entire trial. However, these differences were not found in the ‘medicated’ session. No significant differences were found in pupil size mean or variability in either session. Panagiotidi et al. (2017) found a similar positive association between microsaccade rate and self-assessed ADHD tendencies within a healthy population, but did not investigate pupil size or blink rate.

It is important to note that these studies differ in a significant way. Fried et al. (2014) focused on task-based differences, which arise partly from external circumstances. Healthy controls were able to fixate before target onset, meaning that they were able to control their eye movements to some extent when this was relevant for the task. Those control participants showed a large increase in blinks and microsaccades only after the target has disappeared from the screen. ADHD patients showed deficiencies in this functionality, which was accompanied with decreased task performance. However, Panagiotidi et al. (2017) took a more resting state-based approach, using 20 trials of 20 seconds each, in which participants were asked to fixate on cross, without any additional task or stimuli. This type of paradigm, in which all circumstances are kept equal, captures mostly endogenous variability by default.

It may be tempting to attribute the observed effects to individual differences in ‘attention’. However, in the paradigms of Fried et al. (2014) and Panagiotidi et al. (2017), ‘attention’ may manifest in different ways – the latter relates to internal fluctuations over time, while the former paradigm makes use of covert attention. As described in the above section on *Oculomotor functioning and variability*, these reflect distinct phenomena, and as such, they may not necessarily have similar outcomes.

This is of importance, because ADHD may affect a multiplicity of neuropsychological domains, which means that even when certain behavioural deficiencies or differences are found, it can be hard to pinpoint through which mechanism(s) these arise. While Panagiotidi et al. (2017) did report that the ADHD questionnaire they used is comprised of two subscales – inattention and impulsivity/hyperactivity – analyses were only conducted on the total scores, because both scales correlated highly to the total scores (r-values of .81 and .90 respectively). However, these types of high correlations between subscales and total scores are to be expected, as questionnaires tend to measure one construct, and the total scores reflect nothing more than the sum of the parts. As the correlation between the subscales was only moderate (r = .46), the subscales show sufficient non-shared variance (78.8%) to investigate their separate contributions. Analysing the subscales separately may still reveal potential differences between them, particularly when it is unclear what exact mechanism causes the correlation.

### Current research

In the current research, we examine the resting state paradigm for eye movements in more detail, to see if it produces reliable markers within individuals over different time points (repeatability) and over different conditions (generalisation). In particular, we will be looking at microsaccade rate, pupil size variability, blink rate, and gaze variability (in both horizontal and vertical dimension). We also aim to further explore the relationship of oculomotor variability to self-assessed ADHD symptomatology.

Impulsivity is one of the main characteristics of ADHD, and previous literature has associated self-assessed ADHD tendencies with impulsivity (Berg, Latzman, Bliwise & Lilienfield, 2015; Miller, Derefinko, Lynam, Milich & Fillmore, 2010; although some facets of impulsivity may be more important than others). ADHD has also been associated with increased mind wandering (Shaw & Giambra, 1993; Seli et al., 2015). Shaw and Giambra (1993) furthermore investigated mind wandering in undiagnosed college students and found that participants who scored in the lowest tier of self-assessed ADHD symptoms during childhood were less prone to mind wandering than participants who scored in the highest tier. Possibly, this reflects a decreased tendency to keep top-down focus with increased ADHD tendencies. To get further insight into the mechanisms underlying potential individual differences in oculomotor variability, we therefore also included self-assessed measures of mind wandering and impulsivity.

We aim to replicate positive associations of these two measures with self-assessed ADHD, as well as investigate their relationship to oculomotor variability.

Figure 1 show an overview of our three aims. To examine these aims, we combined data of three behavioural experiments (129 participants in total). In Experiment 1 and 2, participants took part in a four minutes long resting state paradigm and repeated this half an hour to 50 minutes later after they completed a computerised task. In Experiment 3, participants took part in a resting state paradigm in three different conditions repeatedly over four different days, with each resting state being one minute long. In all three experiments, participants filled in questionnaires on ADHD tendencies, mind wandering tendencies, and impulsivity. This allowed us to investigate: 1) the intra-individual reliability of oculomotor behaviour, 2) the between-subject correlations on the questionnaires, and 3) the between-subject correlations between oculomotor behaviour and the questionnaires. Because the predictions for all three questions are highly similar across experiments, they are discussed together below, and analyses were combined whenever possible.

**Figure 1.**
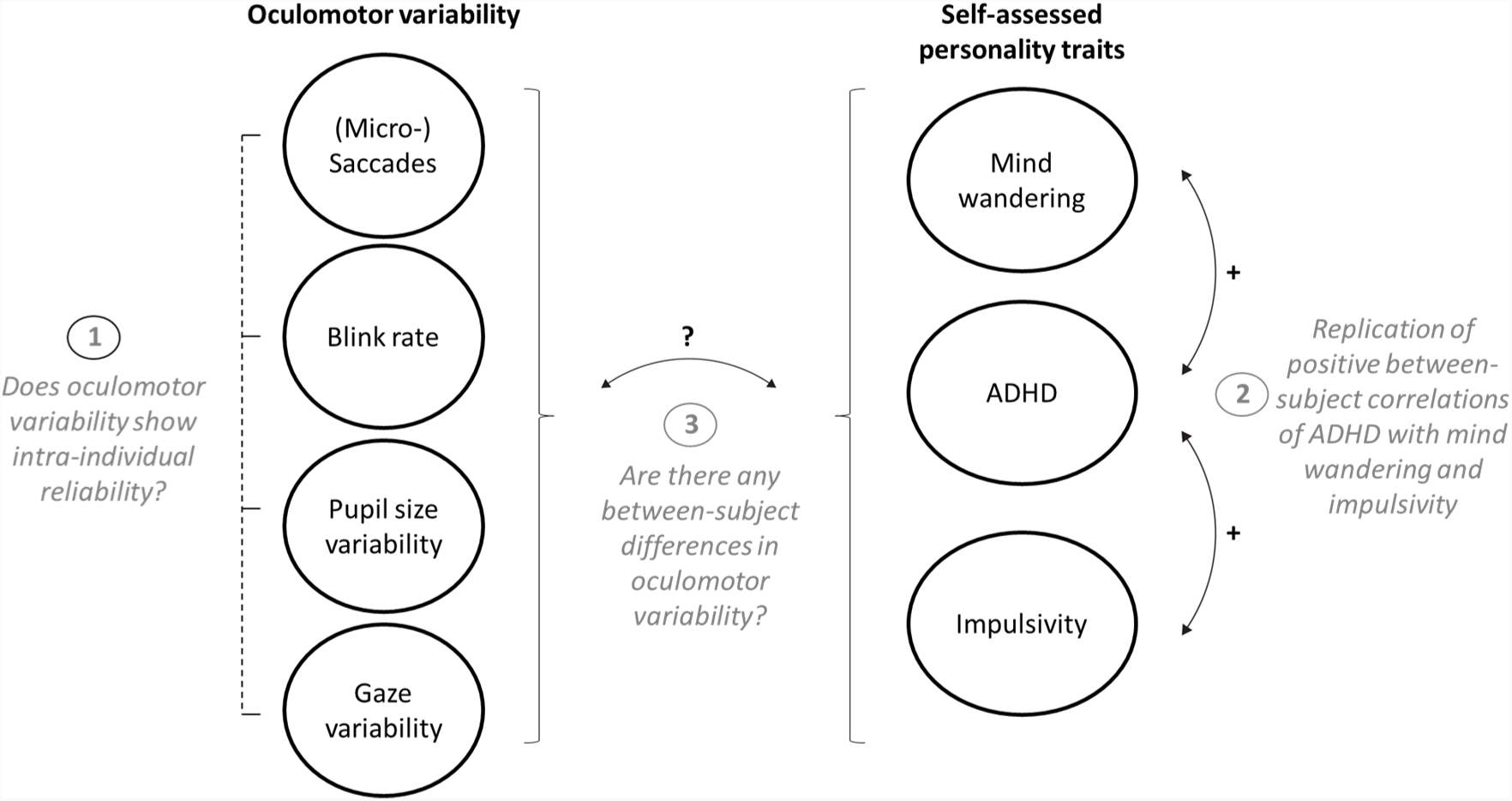
Graphical representation of the oculomotor measures and self-assessed personality traits, with the three aims of the current study.

#### Aim 1. Intra-individual reliability of oculomotor behaviour

If variability of oculomotor functioning is to make a good marker for personality traits, it should show reliability within individuals. We therefore examined the intra-individual reliability of the markers (variability in gaze, pupil size variability, and blink rate in all three experiments, plus microsaccade rate in Experiments 1 and 2) over different points in time on the same day (Experiment 1 and 2) and over different conditions and different days (Experiment 3).

To examine this, mean scores were calculated on each of the different measures for every participant, separately for each resting state (reflecting time/condition). If a measure shows intra-individual reliability, it should correlate highly with itself over the different resting states. Because of the differences in design, the intra-individual reliability was examined separately for each experiment.

#### Aim 2. Between-subject correlations between ADHD, mind wandering, and impulsivity

After completing the resting state paradigms, participants filled in questionnaires on ADHD, mind wandering, and impulsivity. Based on previous literature, we would expect a positive correlation between self-assessed ADHD tendencies and self-assessed mind wandering (Shaw & Giambra, 1993; Seli et al., 2015). Furthermore, we expect a positive correlation between ADHD and impulsivity, similarly based on previous literature (Berg et al., 2015; Miller et al., 2010). Data were combined for all three experiments.

#### Aim 3. Between-subject correlations between questionnaires and oculomotor behaviour

Next, one overall mean was calculated for every participant, separately for each oculomotor measure, collapsed over all time points and conditions. These means were correlated to the ADHD scores, to test if ADHD tendencies are associated with higher oculomotor variability. Furthermore, the scores were correlated to the two subscales of the ADHD questionnaire scores (Inattention and Hyperactivity), as well as to the impulsivity and mind wandering questionnaire scores. The correlations were calculated on the combined data from all three experiments.

If a potential relationship between ADHD and oculomotor variability is caused mainly by a lack of attentional task maintenance, one could expect similar correlations of oculomotor variability to mind wandering and inattention. Alternatively, higher correlations to impulsivity and hyperactivity may reflect that the relationship is driven by a lack of inhibition.

## Methods

### Participants

In total, data of 129 participants was collected. All of them had normal or corrected-to-normal vision. The studies were approved by the local ethics commission.

***Experiment 1.*** Eighty-one participants (66 female, fourteen male, one other, aged between 18-25) contributed in exchange of course credits. Of them, 73 had valid eye tracking data. For three of these remaining 73, the second session was not included because they had more than 33% missing samples.

***Experiment 2.*** Twenty-one participants (eighteen female, 21-40 old, *M*_*age*_ = 26.3) contributed in exchange of a monetary reward. Two of them only took part in one test day, due to technical issues. For another three participants, the second session on the first day was excluded, and for one participant, the second session of the second day was excluded, because more than 33% samples were missing.

***Experiment 3.*** Twenty-eight participants (eighteen female, 18-36 years old, *M*_*age*_ = 25.5) contributed in exchange of a monetary reward, and twenty-six of them had valid eye tracking data. Of these twenty-six participants, one participant had only three sessions, and another had only two sessions. Furthermore, another eleven (out of 303 remaining) sessions from five different participants were not included because more than 33% missing samples were missing.

### Materials

The resting state paradigms were generated with MATLAB (The Mathworks, Inc.) and Psychtoolbox-3 (Brainard, 1997; Kleiner et al., 2007; Pelli, 1997). The background of the paradigms was set at light-grey, and the fixation point was white. An Eyelink 1000 (SR Research) was used in each of the experiments for eye data recording. Each experiment started with calibration and validation with the eye tracker (five-dot calibration in Experiment 1, nine-dot calibration in Experiment 2 and 3). Participants were seated in a chin-rest to limit head movement.

The Adult ADHD Self-Report Scale (ASRS-v1.1; Kessler et al., 2005) was administered to measure ADHD tendencies. The ASRS-v1.1 consists of 18 items with a 5-point scale from 0 (*“Never”*) to 4 (*“Very often”*) and has a high reliability (with Cronbach’s α ranging from .88 to .94; Adler et al., 2006; 2012). The ASRS-v1.1 can be divided into two subscales – Inattention and Hyperactivity / impulsivity -reflecting the two main subtypes of ADHD (Kessler et al., 2005; Reuter, Kirsch & Hennig, 2006).

Furthermore, the Daydreaming Frequency Scale (DFS; Singer & Antrobus, 1963) was administered to measure mind wandering in daily life. The DFS is a subscale of the Imaginal Processes Inventory and measures the amount of daydreaming and off-task mind wandering in daily life. It consists of 12 items, each with a 5-point scale. It has a high internal consistency (Cronbach’s α = .91) and a high test-retest reliability (.76 with an interval of maximum one year; Giambra, 1979-1980).

To measure impulsivity, participants completed the UPPS-P Impulsive Behaviour Scale (Whiteside & Lynam, 2001; Lynam, Smith, Whiteside & Cyders, 2006). The UPPS-P consists of 59 items, with a scale ranging from 1 (*“agree strongly”*) to 4 (*“disagree strongly”*), divided over five subscales: positive urgency, negative urgency, (lack of) premeditation, (lack of) perseverance, and sensation seeking.

#### Experiment 1

The paradigms were generated with a Viglen Genie PC and displayed on an ASUS VG248 monitor with a resolution of 1920 by 1080 and a refresh rate of 144 Hz. Eye movements and pupil dilation were recorded binocularly at 500 Hz.

#### Experiment 2

The resting state paradigms were generated on a HP Z230 Workstation PC and an LG 24GM77 monitor with a resolution of 1920 by 1080 and a refresh rate of 120 Hz. The paradigms were displayed on a projector screen. Eye movements and pupil dilation were recorded binocularly at 500 Hz.

#### Experiment 3

The resting state paradigms were generated with a Bits# Stimulus Processor video-graphic card (Cambridge Research Systems) and a Viglen VIG80S PC, and were displayed on an hp p1230 monitor with a resolution of 1280 by 1024 and a refresh rate of 85Hz. Eye movements and pupil dilation were recorded monocularly at 1000 Hz.

### Design

#### Experiment 1 and 2

Resting state eye movements and pupil dilation were recorded before and after a behavioural task – see Figure 2 for an overview. This gave (2 × 4) 8 minutes of resting state eye measures in total for each participant. ADHD tendencies, mind wandering tendencies, and impulsivity characteristics in daily life were measured with questionnaires.

**Figure 2.**
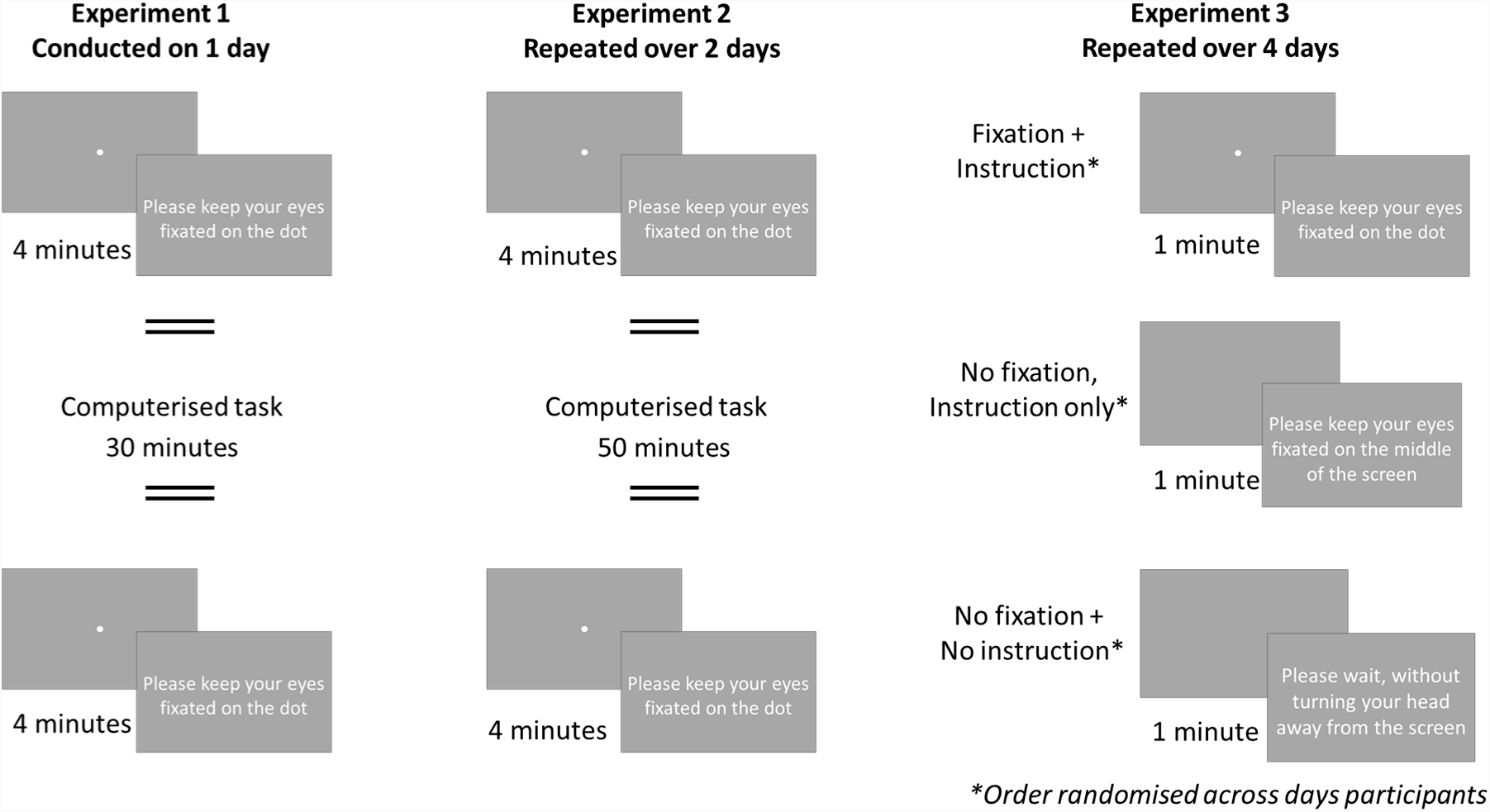
Overview of the resting state eye movement paradigms of all three experiments.

#### Experiment 3

Resting state eye movements and pupil dilation were recorded in three different condition – see Figure 2 for an overview. In the ‘Fixation plus instruction’-condition, participants were asked to fixate on a fixation dot that was displayed on the centre of the screen. In the ‘No fixation, Instruction only’-condition, participants were shown a blank screen, and were asked to fixate on the centre of the screen. In the ‘No fixation plus no instruction’-condition, participants were also shown a blank screen, but they were only asked to not turn away from the screen, with no further fixation-related instructions. This procedure was repeated over four days – resulting in (1 × 3 × 4) 12 minutes of resting state measures for each participant in total. ADHD tendencies, mind wandering tendencies, and impulsivity characteristics in daily life were measured with questionnaires. Again, ADHD tendencies, mind wandering tendencies, and impulsivity characteristics in daily life were measured with questionnaires.

### Procedure

#### Experiment 1

Participants came to the lab for a session of about 1.5 hours. They were seated at a distance of 615 cm from the screen. Eyes were tracked binocularly during the resting state for four minutes (*time 1*). Next, participants performed a computerised task, lasting about 30 minutes (data not analysed in the current paper). Right after finishing this task, the resting state paradigm was conducted again (*time 2*). Lastly, participants filled in nine questionnaires: the DFS, ASRS-v1.1, and UPPS-P, as well as the Beck Anxiety Inventory Second edition (Beck et al., 1993), Beck Depression Inventory Second edition (Beck et al., 1996), Short form Wisconsin Schizotypy scales (Winterstein et al., 2011), Five-facet Mindfulness Questionnaire (Baer, Smith, Hopkins, Krietemeyer & Toney, 2008), Toronto mindfulness scale (Lau et al., 2006), and Positive and Negative Affect Schedule (Watson, Clark & Tellegen, 1988). Only the first three questionnaires were analysed in the current study.

#### Experiment 2

Participants came to the lab for two sessions, each about 1.5 hours. They were seated at a distance of 1185 cm to the screen. Eyes were tracked binocularly for four minutes (*time 1*). Next, they performed a computerised task of about 50 minutes (data not analysed in the current paper), and afterwards they conducted the resting state paradigm again (*time 2*). Lastly, participants filled in the DFS, ASRS-v1.1, and UPPS-P.

#### Experiment 3

The experiment consisted of four sessions of about an hour. Participants were seated at a distance of 104 cm to the screen. Eyes were tracked monocularly in the three different conditions. Each condition lasted 60 seconds. Instructions were shown for two seconds. For each participant, the order of the conditions was random on each of the four sessions. After completing the resting state eye movements paradigm, participants completed a 30 to 45 minutes computerised task (data not analysed in the current paper). On the last day, they filled in the DFS, ASRS-v1.1, and UPPS-P.

### Data preparation and analysis

#### Oculomotor measures

Blinks were defined as missing tracking data, with a maximum of 1000 ms. The total number of blinks throughout each session was counted, and a blink rate per second was subsequently calculated. Pupil size variability was calculated by dividing the standard deviation of the pupil size throughout each session by the mean pupil size – reflecting the coefficient of variation (CV). Gaze variability was calculated separately for the x-and y-screen dimension by calculating the standard deviation of position in degrees throughout the entire session (these standard deviations were not normalised by the mean, as the mean degrees in the middle of the screen is approximately zero). To minimise noise, 20 ms were excluded both before and after missing samples from the calculation of the pupil size and gaze variability.

Binocular microsaccade detection (Experiment 1 and 2 only) was done with the algorithm of Engbert and Kliegl (2003). The λ value was set to five. To reduce noise in the detection process, saccades were defined as being at least three samples long. Furthermore, a period of 100 ms both prior and following blinks was excluded. Missing/excluded samples were subsequently interpolated. To avoid the false detection of post-saccadic oscillations as microsaccades, a window of 20 ms following each saccade was excluded. Saccades with amplitudes above 2° or with peak velocities above 200°/s were excluded from subsequent analyses. To sanity check the microsaccades, saccade amplitude was correlated with velocity over all participants and over both time points (also known as the ‘main sequence’). These were highly correlated to each other for both Experiment 1 (r = .88, BF_10_ = ∞, p < .001) and for Experiment 2 (r = .86, BF_10_ = ∞, p < .001). The mean microsaccade rate was 1.1 per second (SD = .43) for Experiment 1 and 1.58 (SD = 47) for Experiment 2, which is within the typical rate of 1-2 per second (Ciuffreda & Tannen, 1995).

Distributions of the oculomotor measures were highly skewed on the group level. This may bias the results of the correlation analyses, particularly for Experiment 2 and 3, which have smaller sample sizes. For consistency, all analyses were conducted on the natural logarithm of the measures.

#### Questionnaires

Scores on items of the questionnaires were reversed when necessary. Missing responses were substituted with the median (but note that the number of missing responses was neglectable, .26%). Next, the total score was calculated for each of questionnaire. Individual item scores were used to check the questionnaires internal consistency (Cronbach’s α; Cronbach, 1951) – see Table 1 for an overview.

**Table 1.**
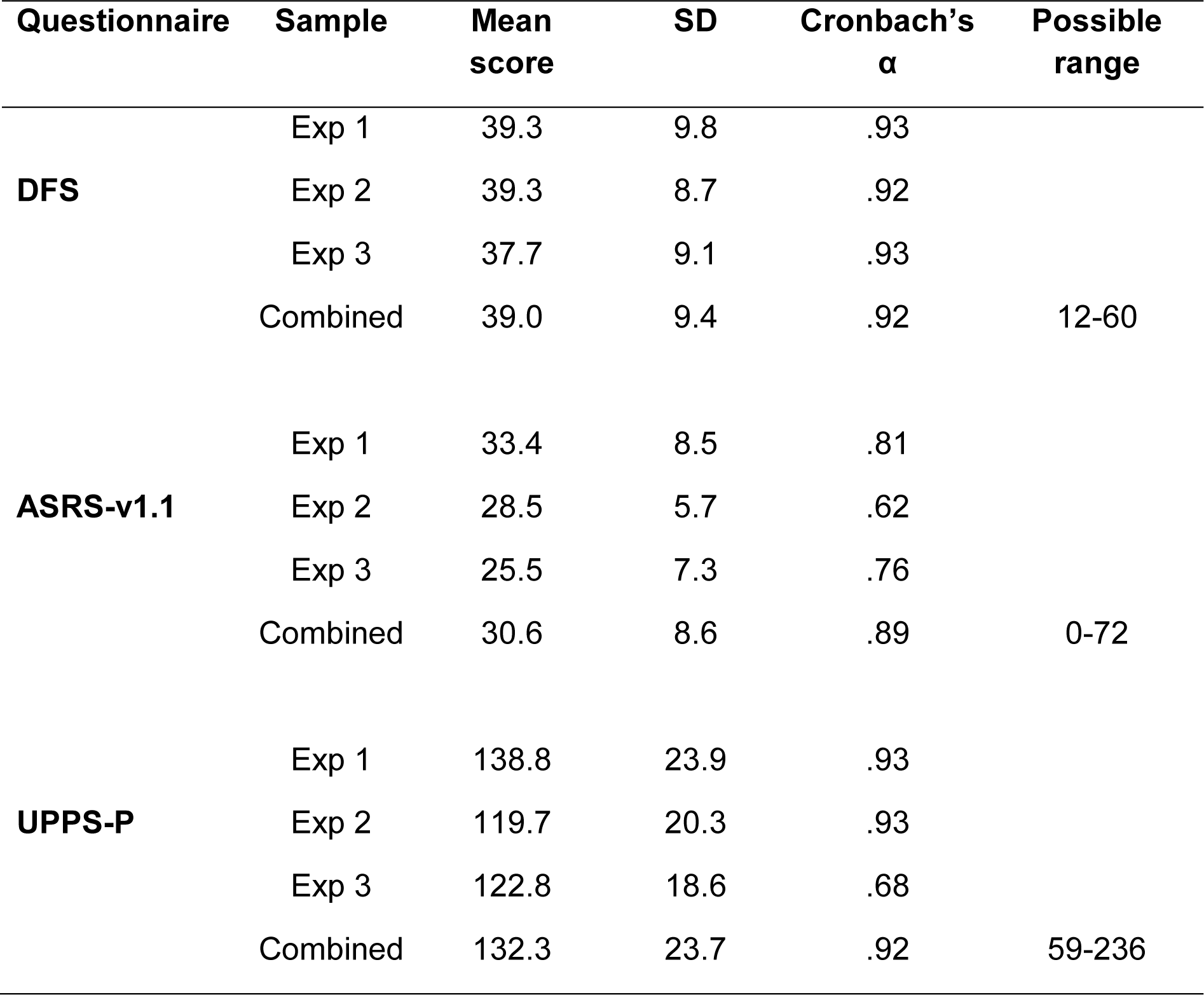
Overview of the Daydreaming Frequency Scale (DFS), the Adult ADHD Self-Report Scale (ASRS-v1.1), and the UPPS-P Impulsive Behaviour Scale (UPPS-P). Shown are the mean scores and standard deviations (SD) over all the participants, as well as the internal consistency (Cronbach’s α) for each questionnaire, for each sample separately as well as for the combined data. Also shown are the minimum and maximum possible scores of each questionnaire.

## Results aim 1. Intra-individual reliability of oculomotor variability measures

### Experiment 1. Reliability over time

Two means were calculated for each measure (microsaccade rate, blink rate, pupil size variability, gaze-x variability, and gaze-y variability): One for time point 1 (pre-task) and one for time point 2 (post-task). Bayesian Pearson pairs were then conducted on each of the measures to test intra-individual reliability over time. Figure 3 shows the within-subject correlational plots over the two time points for the logged measures of gaze variability in the horizontal and vertical dimension, pupil size variability, blink rate, and microsaccade rate – with correlation coefficients and logged Bayes Factors (BF_10_) on top.

**Figure 3.**
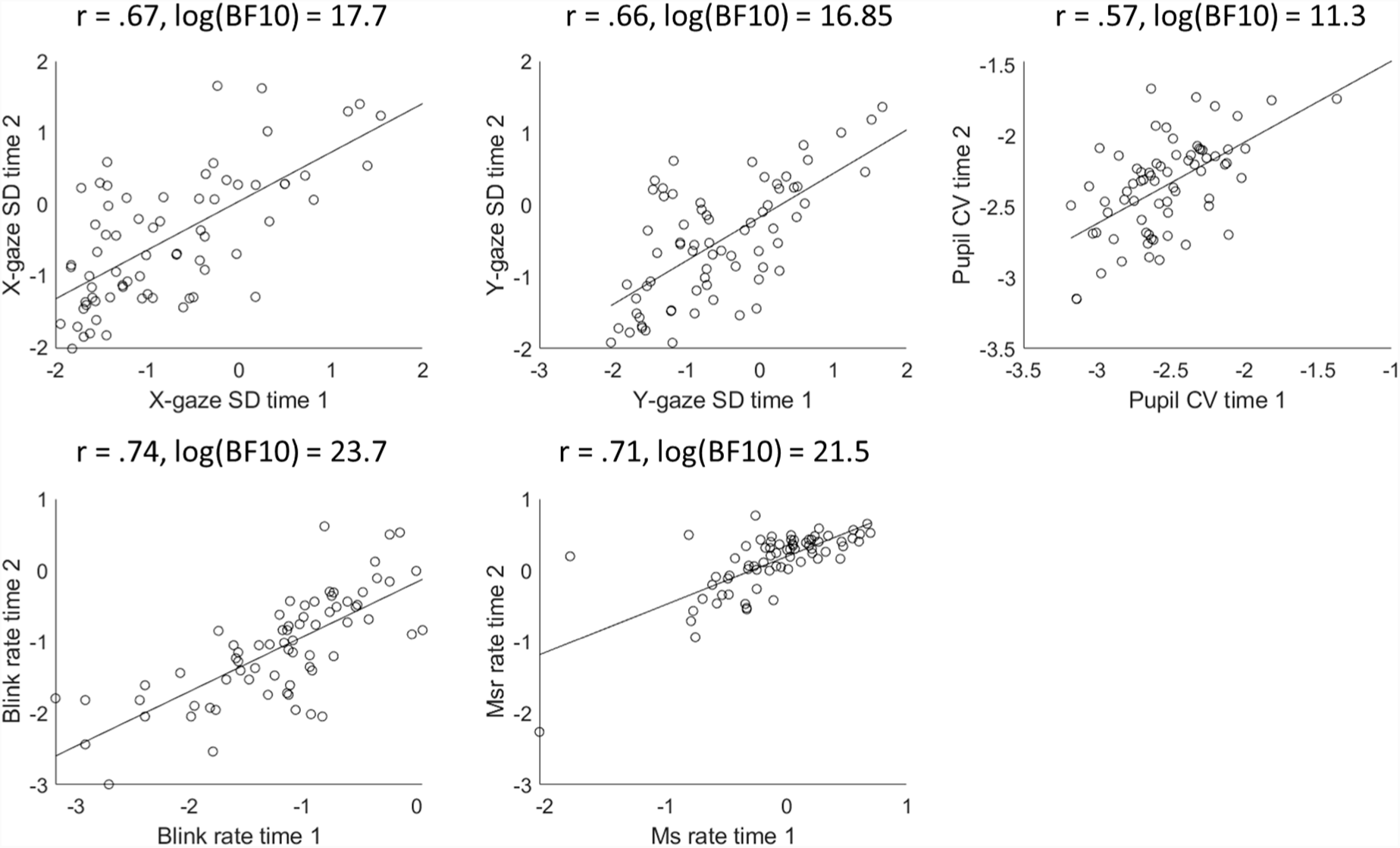
Correlations between time point 1 (pre-task) and time point 2 (post-task) for each of the five oculomotor measures from Experiment 1: Gaze variability (standard deviation; SD) in the horizontal dimension, gaze SD in the vertical dimension, pupil size coefficient of variability (CV), blink rate per second, and microsaccade rate per second (Ms). All five measures show a high correlation coefficient and accompanying high Bayes Factor, indicating that the measures show intra-individual reliability over time. Note that both the measures and the Bayes Factors are logged.

The BF_10_ reflect the likelihood of the data for the alternative hypothesis (in this case, the presence of a correlation) over the null-hypothesis (in this case, the absence of a correlation), and can take a value between zero to infinity.^1^ For example, for gaze variability in the horizontal dimension, the log(BF_10_) between time 1 and 2 is 17.7 – meaning that the likelihood of the data is (exp(17.7) =) 48642102 times larger under the alternative than under the null-hypothesis. This can be interpreted as extremely high evidence for the presence over the absence of a correlation between the two time points. The other four measures show similarly extreme Bayes Factors. Each of the measures show high and positive r-values, indicating that they show intra-individual consistency. Thus, oculomotor shows reliability when measured half an hour apart.

### Experiment 2. Reliability over time and days

Combined, Experiments 2 and 3 have 21 correlation pairs for each oculomotor measure, each testing the reliability over different time points and days. Rather than having to plot each correlation separately and then trying to assess the global patterns, the distributions of these correlations are shown in violin plots (Figure 4). This way of representing the data allows for an immediate overall picture of the correlations. The vertical dimension of these violin plots indicates the entire range of correlation coefficients (top panel) and accompanying Bayes Factors (bottom panel), while the horizontal dimension indicates the density. Each condition is also plotted (coloured triangles and asterisks), with the white dot representing the median value.

**Figure 4.**
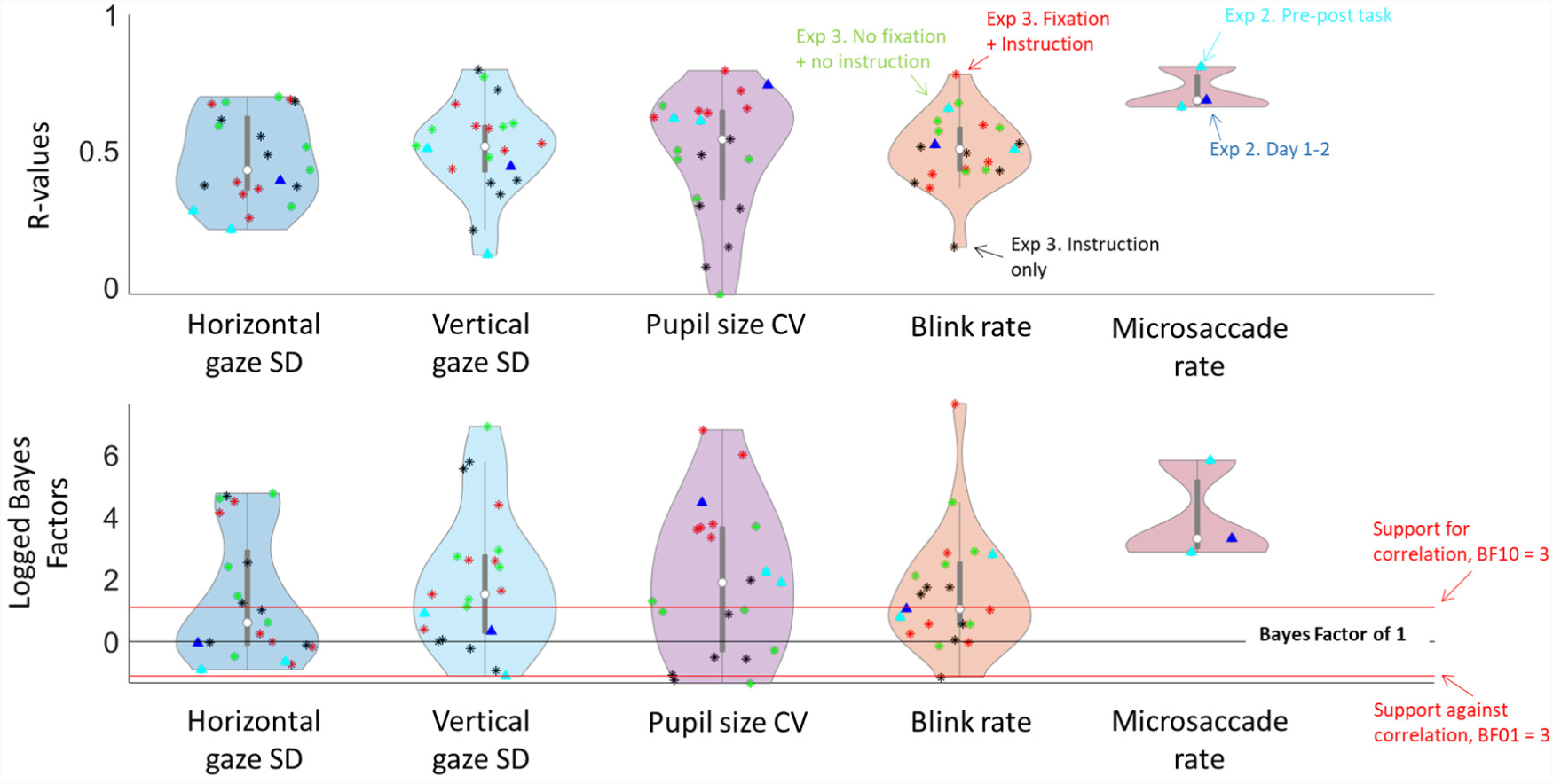
Distributions of the correlation coefficients (top panel) and accompanying logged Bayes Factors (bottom panel) of the correlation analyses on within-subject reliability for each of the five oculomotor measures. Values denoted with a triangle represent the correlations for Experiment 2, with light-blue triangles representing the correlations between different time points (pre and post task), and dark-blue triangles representing the correlation between days. Values denoted with an asterisk represent the correlations for Experiment 3, with red, black, and green representing the different conditions (‘Fixation plus instruction’, ‘No fixation, instruction only’, and ‘No fixation plus no instruction’ respectively). In the top panel, higher values on the y-axis indicate higher correlation coefficients. In the bottom panel, values above the upper red line indicate evidence in favour of the existence of correlations over time, while values below the lower red line (log(BF) < −1) indicate evidence against correlation over time. Values falling between the two red lines are interpreted as indeterminate. Overall, reliability seems low for variability in gaze position, particularly in the horizontal dimension, but the other measures show good reliability.

To test the intra-individual reliability over time in Experiment 2, four means were calculated for each measure (microsaccade rate, blink rate, pupil size variability, gaze-x variability, and gaze-y variability): One for time 1 (pre-task) and one for time 2 (post-task), both for day 1 and day 2. For both days, Bayesian Pearson pairs were conducted between time 1 and time 2 on each measure – giving two replications of the analysis of Experiment 1 (shown in Figure 4 in light-blue triangles). Again, we found evidence in favour of correlations between time 1 and 2 for pupil size variability, blink rate, and microsaccade rate (with all six BF_10_ above 1, and only one of them in the indeterminate range), with corresponding r-values all being moderate to high. These findings again indicate good intra-individual reliability of the measures – especially when considering the much smaller sample size of this experiment. These results replicate the findings from Experiment 1 with almost twice as much time in between the two time points. However, we no longer found evidence for intra-individual reliability in gaze variability, especially in the horizontal dimension: All four BF_10_ were in the indeterminate range, with three of them being below 1.

Next, means over time points were averaged, resulting in two means for each measure: One for day 1, and one for day 2. Bayesian Pearson pairs were conducted on each of the measures between day 1 and day 2 to test intra-individual reliability on a longer time-span. Figure 4 shows the correlation coefficient and Bayes Factor for each measure (dark-blue triangles). The correlations between days show similar patterns to the ones between time points: Gaze variability appears least reliable, while pupil size variability, blink rate, and microsaccade rate show good reliability.

### Interim-discussion: How long should a resting state session be?

Overall, oculomotor variability showed good intra-individual reliability over time, both before and after a task of 30/50 minutes (Experiment 1 and 2 respectively), as well as over days (Experiment 2) – although variability in gaze position appeared to be the least reliable measure. It should be noted that the differences we found between individuals are substantial – for example, in Experiment 1, for gaze variability in the horizontal dimension at time 1, the most variable participant has an SD that is 32 times larger than the least variable participant.

Findings for both experiments were based on a resting state of four minutes. The next question may be how long a resting state session should minimally take before it could be considered to produce reliable measures. To answer this question, we analysed the data of Experiment 1 – looking at variability in gaze and in pupil size over the course of the resting state. First, for each measure, the Pearson r-value between time 1 and time 2 was calculated on every cumulative second. This results in 240 r-values – with the first r-value being based on one second of data, and the last r-value being based on four minutes of data. This trajectory reflects how the consistency between the two time points develops as more data is collected (red line on Figure 5).

**Figure 5.**
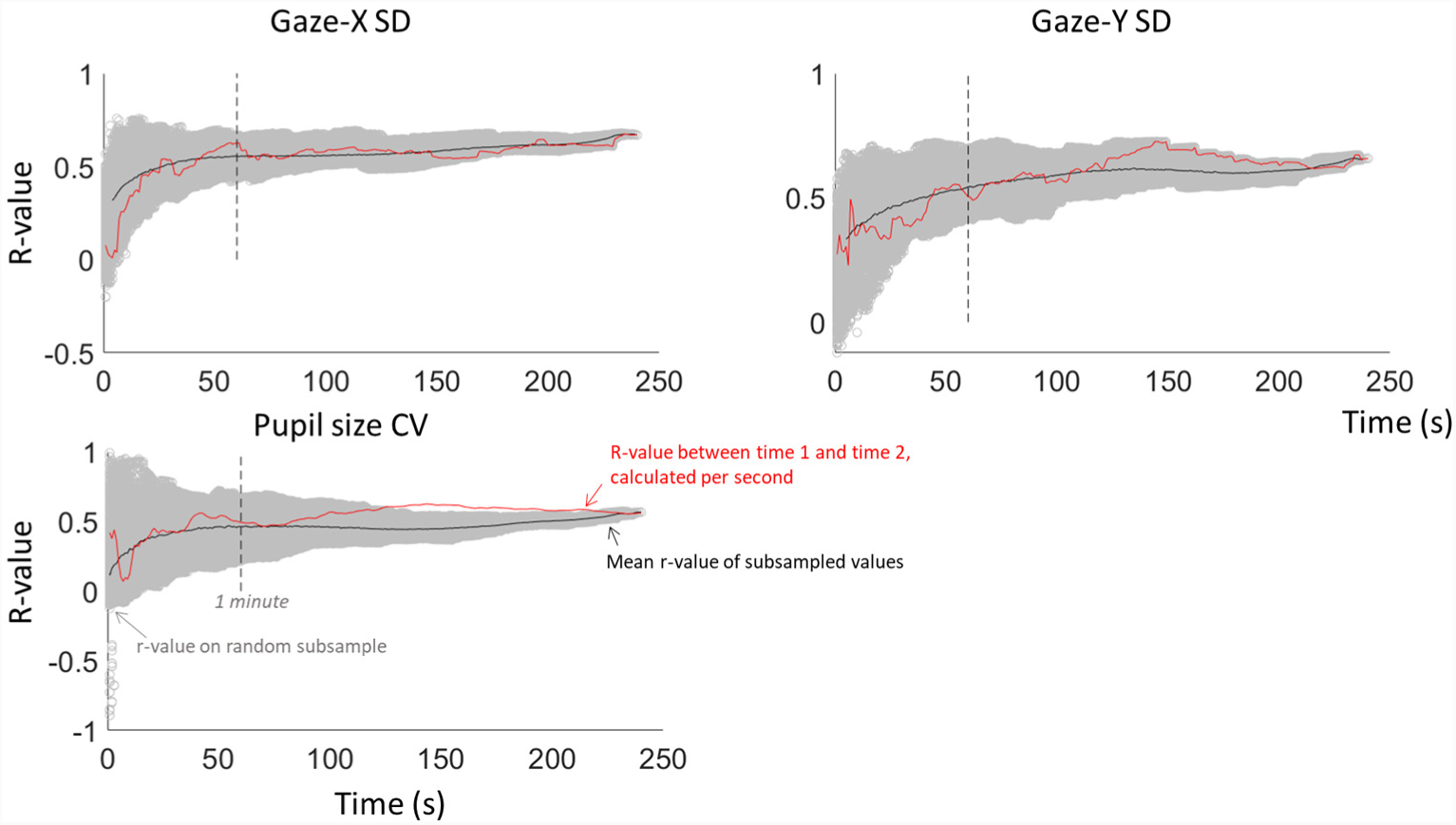
Intra-individual reliability of Experiment 1 over the course of the resting state for our three continuous measures: gaze variability (in the horizontal and vertical dimension) and pupil size variability. The r-value between time points 1 and 2 was calculated at each cumulative second (red), thus reflecting the trajectory over time. Next, for each cumulative second, estimates of the r-value were calculated on 1000 random subsamples. These estimates are shown in light-grey circles, with the mean of these subsamples shown in black.

Next, we adopted a subsampling approach, using a simplified version of Schonbrodt and Perugini’s (2013) approach. From the entire pool of data of four minutes, one chunk of data was randomly selected for both time points, and the r-value between them was calculated. This subsampling was done 1000 times for each cumulative second, represented on Figure 5 by the grey circles, with the mean represented by the black line. This means that, for example, at time = 1 sec, there are 1000 different r-values, each based on one continuous randomly selected second in the entire pool of data. Next, at time = 2 sec, there are also 1000 different r-values, each based on two continuous randomly selected seconds in the data. As such, we end up with 1000 r-values at each cumulative second. Because of this method, the r-values converge to one point as the subsamples are based on more data – resulting in very small margins of error at the right side of the x-axis. Still, the mean trajectory of the subsampled r-values combined with the trajectory of the ‘actual’ r-values can give an idea of the minimal necessary length for an oculomotor resting state.

Looking at Figure 5, it seems that reliability is lower and more volatile when it is based on less than a minute of data. After one minute, the reliability stabilises, and does not seem to improve any further after two minutes. Based on these outcomes, we recommend that an oculomotor resting state session is no shorter than one minute, but that it may not be necessary to collect more than two minutes of continuous data.

In Experiment 3, we were not only interested in the intra-individual reliability of oculomotor variability over different days (repeatability), but also in the extent to which the oculomotor variability would generalise over different types of ‘oculomotor resting states’. For this, we used the same resting state version as in Experiment 1 and 2, as well as a free viewing version (in which participants did not have to fixate on anything, and were free to look anywhere on the screen), and an ‘intermediate’ version (in which participants were asked to fixate on the middle of the screen, but were not provided with a fixation dot). Because participants were asked to participate in each condition on four different days (resulting in twelve resting states per participant), we made the sessions shorter – using one minute per resting state instead of four. As shown above, this is long enough to produce reliable estimates.

### Experiment 3. Reliability over days and conditions

For each of the measures (blink rate, pupil size variability, horizontal gaze variability, and vertical gaze variability), means were calculated separately for each condition and each day (thus resulting in twelve means for each measure). Bayesian Pearson correlations were conducted for each measure between the means over the different days, separately for each condition (resulting in eighteen correlation pairs for each) – to test the reliability of the oculomotor measures over time. Figure 4 shows these correlation coefficients and Bayes Factors (asterisks) for each of the three conditions (with ‘Fixation plus instruction’ in red, ‘No fixation, instruction only’ in black, and ‘No fixation plus no instruction’ in light-green). The overall pattern is similar to that of Experiment 2. Gaze variability in the horizontal dimension seems least reliable: Bayes Factors mostly show indeterminate evidence *against* a correlation. Pupil size variability and blink rate show the most evidence for good reliability: While the Bayes Factors show a very wide range (with some below 1, but others logged values around 6-7), the overall distribution favours the existence of correlations over the absence of correlations. Again, correlation coefficients for these two measures were mostly moderate to high, with both median values around .5. Our ‘intermediate’ condition, in which participants were asked to fixate at the middle of a blank screen, appeared to produce the least reliable measures.

Over all three experiments, we thus found reliability in oculomotor measures over time, from relatively short ranges (30 to 50 minutes) up to multiple days apart. Next, we were interested in to what extent the oculomotor measures were generalisable over different types of resting states. To examine this, means were averaged over days, resulting in three means for each measure, each reflecting one condition. Bayesian Pearson correlations were conducted on the means of the three conditions – to investigate the reliability of the measures over different conditions. Figure 6 shows the correlation plots between the conditions for each measure, with Table 2 showing the accompanying correlation coefficients and Bayes Factors. All correlations had a Bayes Factor above 1, with eight of them ranging from moderate to extreme. Overall, the measures again show moderate to high reliability, although it is the poorest for gaze variability in the horizontal dimension. Blink rate seems to have the highest reliability over conditions.

**Table 2.** Overview of the intra-individual reliability across conditions for each of the four measures from Experiment 3. For each pair of conditions and each measure, the correlation coefficient is shown, with the accompanying BF_10_ in brackets.

| Measure | Fixation + Instruction<br>vs<br>Instruction only | Fixation + Instruction<br>vs<br>No fixation + Instruction | Instruction only<br>vs<br>No fixation + Instruction |
| --- | --- | --- | --- |
| **Gaze-X** | .36 (1.12) | .63 (60.05) | .37 (1.28) |
| **Gaze-Y** | .47 (3.66) | .73 (1241.77) | .45 (3.10) |
| **Pupil size** | .40 (1.67) | .83 (77689) | .43 (2.41) |
| **Blink rate** | .84 (241807) | .85 (288824) | .79 (10224) |

**Figure 6.**
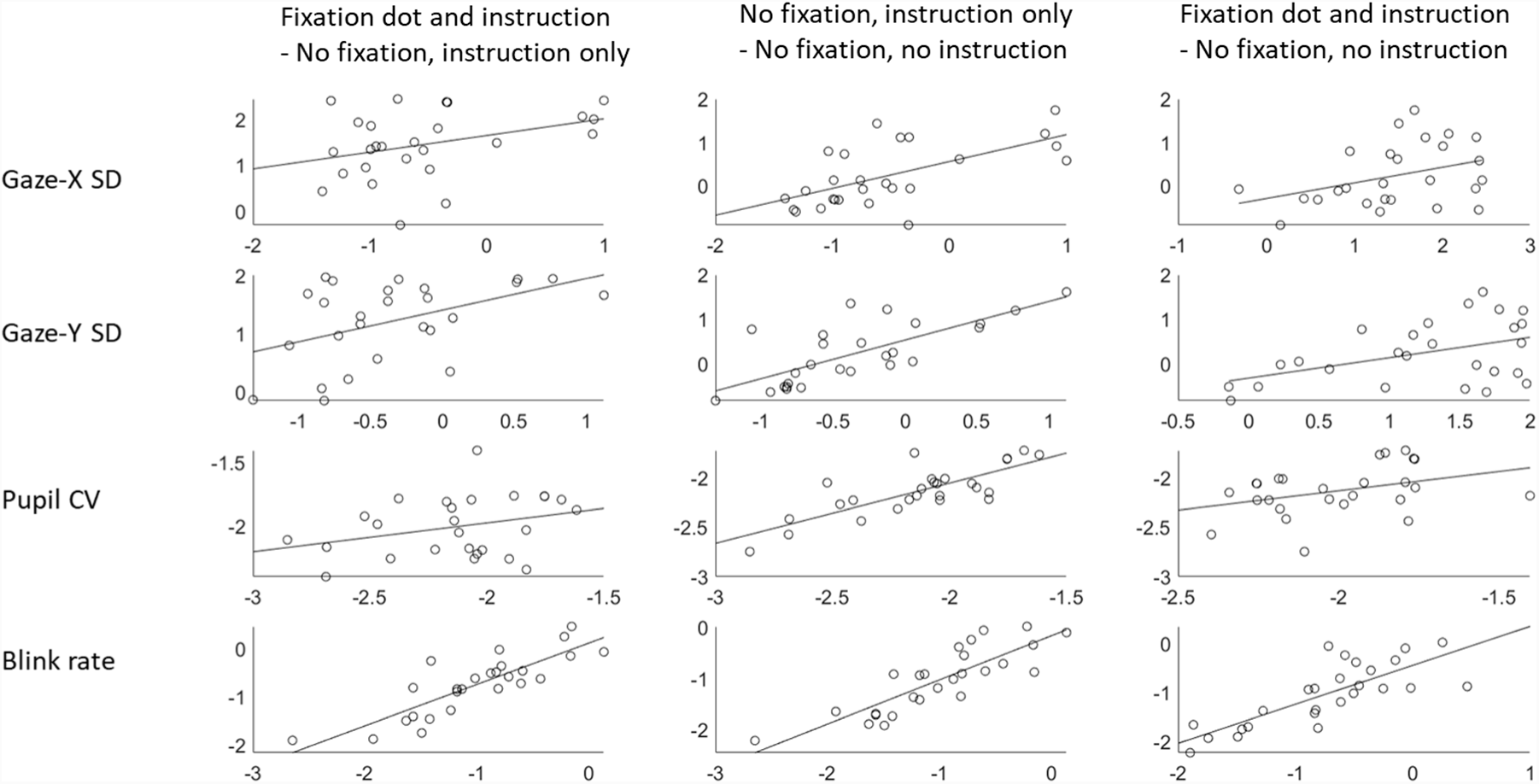
Correlation plots between the three different conditions (‘Fixation plus instruction’, ‘No fixation, instruction only’, and ‘No fixation plus no instruction’) on each of the four oculomotor measures from Experiment 3. Overall, evidence favours the existence of correlations – suggesting good intra-individual reliability of oculomotor variability over the different conditions. Note that the measures are logged.

**Figure 7.**
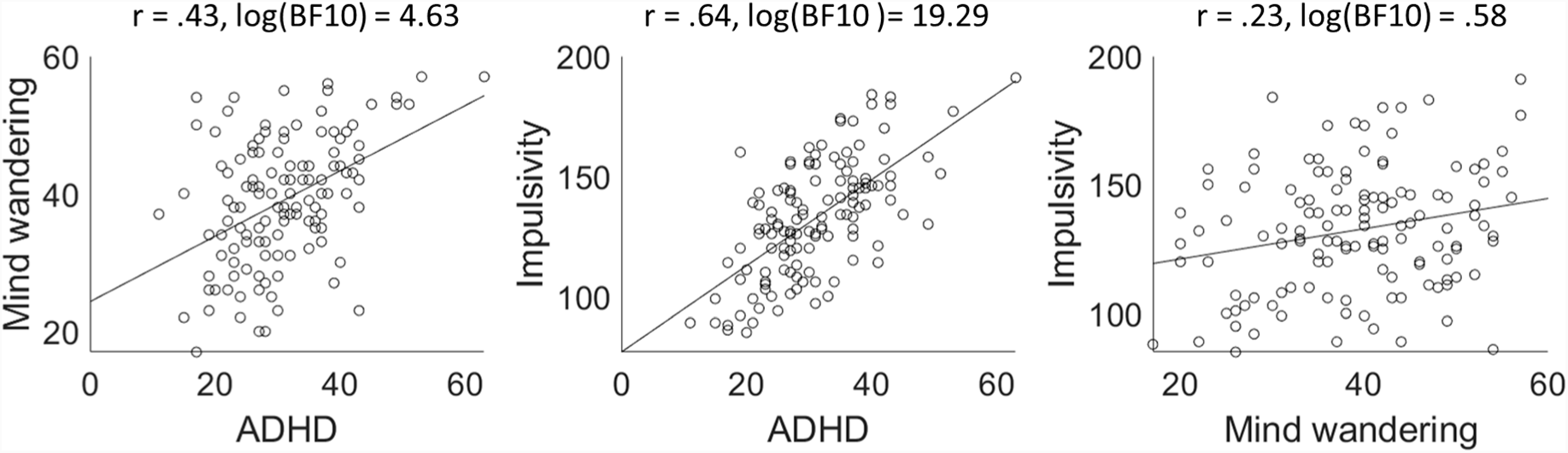
Correlational plots between self-assessed ADHD tendencies, mind wandering tendencies, and impulsivity, with accompanying Pearson r and Bayes Factor values. ADHD tendencies are positively correlated to both mind wandering and impulsivity – replicating previous literature.

**Figure 8.**
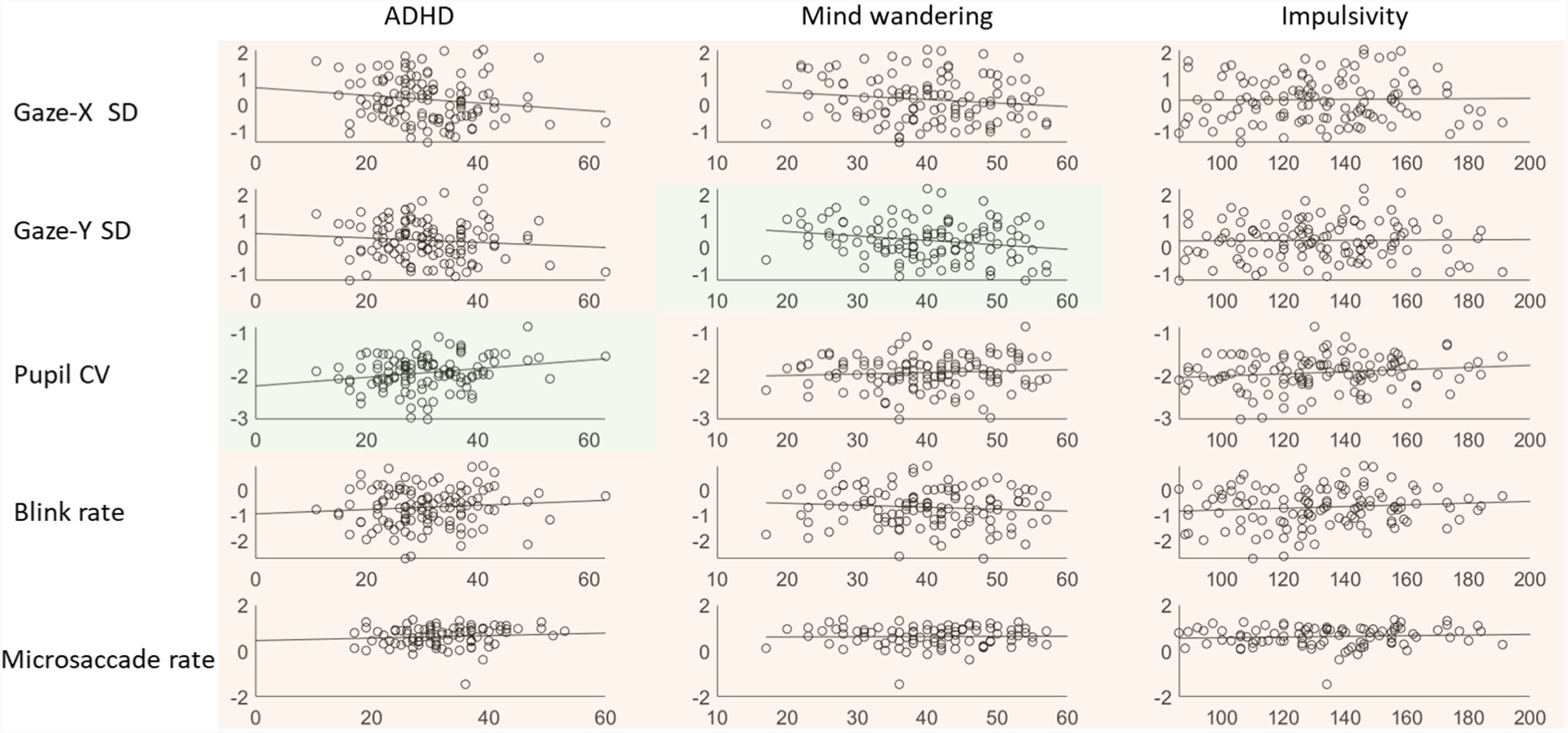
Correlation plots between the oculomotor measures and the self-assessed personality traits. Green shading indicates that the corresponding Bayes Factor is above 1 (indicating evidence in favour of a correlation between the conditions on that measure), while red shading indicates a Bayes Factor below 1 (indicating evidence against a correlation). Note that the oculomotor measures are logged.

#### Intra-class correlation

The intra-class correlation can estimate the reliability of a larger group of measures, to reflect to what extent they measure the same underlying phenomenon – and as such, can reflect the ‘correlation’ between more than two measures. To estimate the intra-class correlation, a two-way random model was conducted on each measure. The measure of consistency was estimated, as this is most similar to our Pearson correlation analyses. Table 3 shows the correlation coefficients for the average measure, to reflect the overall consistency of the resting states. The analysis was run both on each condition separately as well, to get an estimate of reliability over days, and collapsed over conditions and days, to get an estimate of the overall reliability of the paradigm.

**Table 3.**
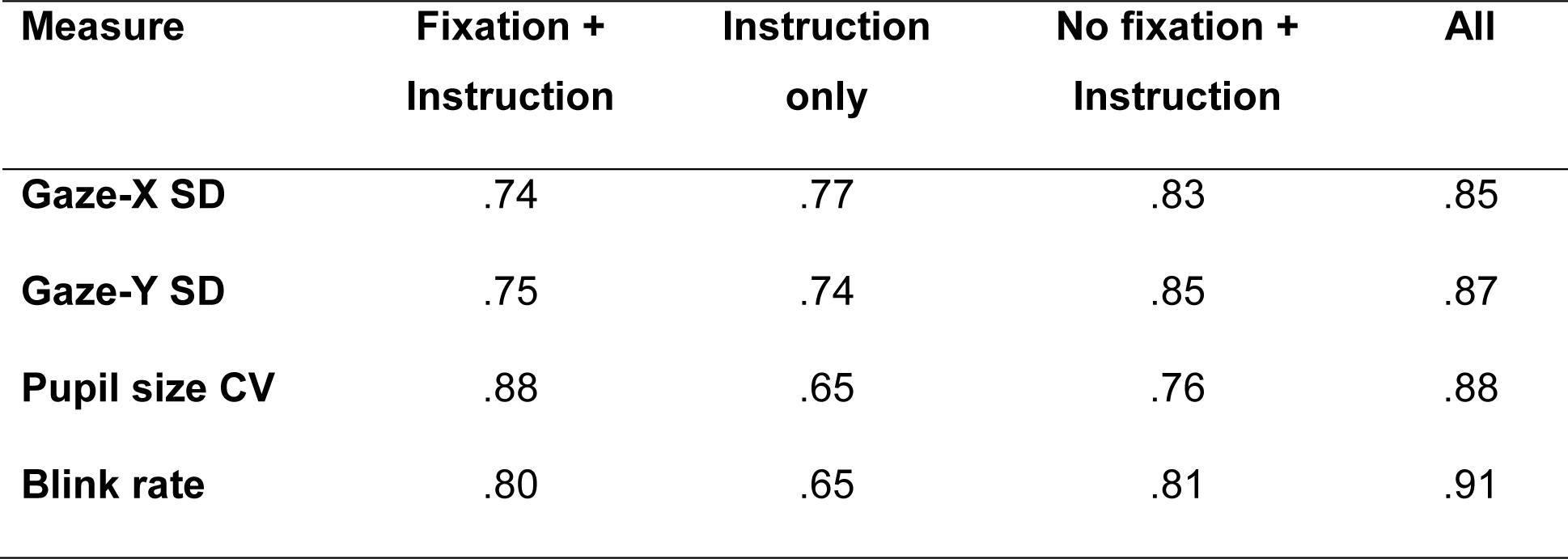
Overview of the intra-class correlation coefficients of the average measure for each of the three conditions from Experiment 3, separately for each of the four measures, as well as the coefficients per measure over all conditions and days combined.

| Measure | Fixation +<br>Instruction | Instruction<br>only | No fixation +<br>Instruction | All |
| --- | --- | --- | --- | --- |
| Gaze-X SD | .74 | .77 | .83 | .85 |
| Gaze-Y SD | .75 | .74 | .85 | .87 |
| Pupil size CV | .88 | .65 | .76 | .88 |
| Blink rate | .80 | .65 | .81 | .91 |

**Table 4.** Pearson r-values (BF10) between the three questionnaires and the measures of oculomotor variability, combined over the three experiments.

| Measure | ADHD | Mind wandering | Impulsivity |
| --- | --- | --- | --- |
| **Gaze-X SD** | -.15 (.41) | -.15 (.38) | .02 (.12) |
| **Gaze-Y SD** | -.10 (.20) | -.21 (1.61) | .02 (.12) |
| **Pupil size SD** | .22 (2.11) | .08 (.16) | .15 (.43) |
| **Blink rate** | .11 (.24) | -.09 (.18) | .12 (.27) |
| **Microsaccade rate** | .10 (.21) | -.02 (.13) | .08 (.17) |

All three conditions showed moderate (.5-.75) to good (.75-.9) reliability (see Koo & Li, 2016 for guidelines), although results again indicate that the ‘Instruction only’ condition produces the least reliable results. When collapsing over all days and all conditions, reliability is even higher, ranging from good to excellent (.9-1). Overall, the conditions seem to measure the same underlying construct – reflecting good intra-individual reliability of oculomotor measures. Interestingly, the coefficients are all at least in the good range, even variability in gaze position – as such, diverging from the results of the individual Pearson correlations. However, the Pearson correlations can only reflect the consistency between two single measures, while our intra-class correlations reflect the consistency over all the different days averaged together. This suggests that over all the days combined, the oculomotor variability still shows within-subject consistency.

## Results aim 2. Between-subject correlations between ADHD, mind wandering, and impulsivity

Bayesian Person correlations were conducted on the questionnaire scores. Figure 6 shows the between-subject correlational plots with their corresponding Pearson r coefficients and Bayes Factors. Looking at the between-subject correlations between ADHD tendencies, mind wandering (DFS), and impulsivity (UPPS-P), we found that ADHD tendencies were highly correlated to impulsivity and mind wandering tendencies. Both of these findings thus provide extreme evidence for replication of previous literature.

There was also some evidence for a correlation between mind wandering and impulsivity, but the evidence was in a much lower range and the accompanying correlation coefficient was similarly low, Pearson r = .23, BF_10_ = 3.8. It seems plausible that this correlation is caused by a confounding effect of ADHD tendencies. To statistically control for ADHD tendencies, a Bayesian Linear Regression was performed in which impulsivity scores were regressed on mind wandering tendencies (alternative Model M_1_) and compared to a null-model that included the ADHD tendencies as model term (model M_0_; see Wetzels & Wagenmakers, 2012 for more details on this method). Bayesian evidence favoured M_0_ over M_1_, BF_01_ = 7.7, indicating that the relationship between impulsivity and mind wandering disappears when controlling for ADHD tendencies.

## Results aim 3. No between-subject correlations between questionnaires and oculomotor behaviour

For each participant, one mean was calculated for each of the measures, collapsed over all potential points of time, days, and conditions. Bayesian Pearson correlations were conducted between these oculomotor measures and the questionnaire scores. Out of the fifteen analyses, ten showed moderate evidence *against* a correlation, and five were in the indeterminate range (three of them with BF_10_ < 1, and the other two with BF_10_ > 1). Looking at the two correlations that had a BF_10_ > 1 (though in the indeterminate range), the accompanying r-values were low (explaining only 4.4 and 4.8% of the total variance).

To examine if any correlations would be more pronounced when looking at the subscales instead of the total scores of ADHD, the inattention and impulsivity/hyperactivity scores were correlated with the oculomotor measures. Pupil size variability correlated with the inattention subscale (r = .24, BF10 = 3.75), but not with impulsivity/hyperactivity (r = .13, BF10 = .31) – indicating that participants with more inattention-related ADHD tendencies showed more variability in pupil size. However, the explained variance was again low (5.8%).

## Discussion

In the current research, we aimed to: 1) examine to what extent endogenous oculomotor variability constitutes a reliable individual trait, 2) replicate positive associations of self-assessed ADHD tendencies to mind wandering and impulsivity, and 3) investigate potential relationships between personality traits and endogenous oculomotor variability. We combined datasets from three experiments including ‘oculomotor resting states’ as well as a set of questionnaires. We found that oculomotor variability indeed shows consistency within individuals, both over time (repeatability) and over different conditions (generalisation). Of the five measures that we used (variability in both horizontal and vertical dimension, pupil size variability, blink rate, and microsaccade rate), each showed consistency to some extent – with blink and microsaccade rate appearing to be the most consistent measures, and gaze variability (particularly in the horizontal dimension) being the weakest.

We also found positive correlations between the self-assessed personality traits, replicating previous associations of ADHD to mind wandering (Shaw & Giambra, 1993; Seli et al., 2015) and impulsivity (Berg et al., 2015; Miller et al., 2010). However, these personality traits did not show convincing correlations with oculomotor variability. Overall, we mostly found Bayesian evidence against correlations – and for the few correlations that were weakly supported, the effect sizes were small. This suggests that within our sample of healthy participants, oculomotor variability did not prove a useful measure for corroborating self-assessed personality traits.

### Reliability of oculomotor variability

Recently, the World Federation of Societies of Biological Psychiatry and the World Federation of ADHD have identified the need for dedicated biomarkers of ADHD (Thome et al., 2012). Oculomotor measures seem appealing: They are easily accessible in terms of money and time, in sharp contrast with typical neuroimaging methods. Indeed, endogenous oculomotor variability has been proposed as a potential biomarker for ADHD (Panagiotidi et al., 2017). However, it is crucial for any biomarker to show intra-individual reliability (Mayeux, 2004).

Intra-individual stability of oculomotor variability during task has been shown in previous research (Andrews & Coppola, 1999; Boot et al., 2009; Castelhano & Henderson, 2008; Poynter et al., 2013; Rayner et al., 2007). Furthermore, there is evidence that oculomotor variability in viewing tasks shows intra-individual correlations with oculomotor variability in the absence of any visual stimulation (‘dark room condition’; Andrews & Coppola, 1999). Because oculomotor variability is measured within tasks, it reflects a mixture of exogenous and endogenous variability. This means that findings can be (partly) driven by individual differences in information processing and strategies. Previous studies have found similar intra-individual reliabilities in reaction time variability across time during task and across different tasks (Hultsch et al., 2002; Saville et al., 2011; Saville et al., 2012; but see Salthouse, 2012). In these contexts, it appears difficult to exactly quantify which part of the variability arises due to exogenous variability, and which part arises due to endogenous variability. To our knowledge, our design is the first to investigate the intra-individual stability in ‘pure’ endogenous variability in oculomotor behaviour – captured by continuous measurement under an absence of changes in the external environment.

When comparing the reliability over time, correlation coefficients (and accompanying Bayesian evidence) were highest in Experiment 1 – in which the two measures were closest together in time – and lowest in Experiment 3 – in which measures were typically separated by multiple days. Still, correlation coefficients showed reasonable intra-individual consistency even in Experiment 3. Of course, the individual correlation pairs will be affected by chance. This is evidenced by the distribution plots in Figure 5, that shows the range of found correlation coefficients is large. However, overall, the distributions favoured moderate to high correlations, with median r-values being around .5 (with the exception of gaze variability). Furthermore, intra-class correlation coefficients showed good to excellent consistency for each of the measures – revealing that, overall, the measures over days appear to measure the same underlying construct. Based on our subsampling analysis on the data of Experiment 1, we can recommend that these sessions should be between 1-2 minutes long, with longer recording sessions being only necessary when the sample is small.

Both over time and over conditions, we found that gaze variability was consistently the weakest measure, particularly in the horizontal dimension. One possibility is that gaze variability is driven by a multitude of sources, including saccades, drift, and tremor, but also by phenomena such as partial blinks. Because of this, gaze variability may have less specificity than the other measures, and thus, less validity as a measure. While reliability and validity are theoretically different constructs, in practice, they often go hand in hand.

It is important to note that our findings show that oculomotor behaviour is consistent within individuals over time – likely reflecting individual traits. This means that individuals who are highly variable at time 1 typically are also highly variable at time 2, while individuals who show low variability at time 1 typically also have low variability at time 2. However, this does not mean that the measures are exactly the same at time 1 and time 2; they are still subject to variability. For example, looking at the reliability over time in Experiment 2 and 3, we typically explain 25% of the total variance (and for Experiment 1, explained variance ranged from 32 to 55%).

### Statistical power and sample size

By combining data from multiple experiments, we were able to study individual differences in oculomotor variability in a large sample. Still, a number of our Bayesian analyses on individual differences produced Bayes Factors in the indeterminate range. If anything, this highlights the importance of using large samples in these types of individual differences studies, especially when considering that effect sizes will typically be small (see Gignac & Szodorai, 2016 for a meta-analysis of effect sizes in Psychology). This may be more apparent with Bayesian analyses, in which evidence is gradual rather than a binary ‘significant versus non-significant’ decision. However, it is also important in traditional ‘null hypothesis significance testing’ (NHST), especially when considering that running underpowered studies actually increases the chance of making Type I errors (Bakker, van Dijk & Wicherts, 2012), while simultaneously making it difficult to interpret non-significant results.

It has been known for a long time that psychological research studies have been largely underpowered due to the use of small sample sizes (Cohen, 1962; Sedlmeier & Gigerenzer, 1989), but that sample sizes have not increased (although this seems partly field-dependent, see Marszalek, Barber, Kohlhart & Holmes, 2011). Researchers in Psychology appear to have incorrect intuitions about statistical power and sample sizes, and rely on rules of thumb and on (incorrect) practices from the literature (Bakker, Hartgerink, Wichters & van der Maas, 2016). To give an example, to obtain the traditionally recommended power of 80% for a correlation with r-value of .3 in NHST, a power analysis shows that a sample of minimally 85 participants is required. In the last years, the topic has gained increasing traction (e.g., Bakker et al., 2012; Button et al., 2013). It should be noted that increasing sample size is not the most obvious and necessary step in all types of studies (Rouder & Haaf, 2018; Smith & Little, 2018), but appears key when studying associations between measures that are prone to large heterogeneity.

However, although the power analysis is a common framework to think about power, sample size is not the only determinant of statistical power (Asendorpf et al., 2013; McCelland, 2000). Among others, one can obtain higher power by testing hypotheses that are well-grounded in theory, avoiding redundancy in predictor variables, increasing differences in experimental conditions, minimising measurement noise during data collection, and using appropriate statistical analyses. The ratio of ‘true variance’ to error variance can further be improved by using reliable measurements, and by collecting enough data points. For instance, using a high-quality eye tracker with a fast refresh rate leads to better oculomotor data, and subsequently to better estimates and higher statistical power.

Our results on individual differences diverge from previous literature, which found a positive association between ADHD and microsaccade rate in a healthy population (Panagiotidi et al., 2017). Comparing our study with theirs, we used the same eye tracker system and refresh rate, as well as the same microsaccade detection algorithm (Engbert & Kliegl, 2003) and the same analysis (Pearson r correlation). However, our study had a higher sample size (our correlation between ADHD tendencies and microsaccade rate included 94 participants, compared to 38 in theirs), as well as more data points (a minimum of 8 minutes in ours compared to ~6.5 minutes). This means that the absence of a replication in our results is not caused by a lack of power.

### Individual differences in oculomotor variability

Potentially, these different findings may be explained by differences in design. In our experiments, oculomotor variability was recorded over a continuous ‘resting state’ session, while in Panagiotidi et al. (2017) participants were asked to fixate for only 20 seconds in a row over 20 different trials. After each trial, they were given a break, and could decide themselves when to continue with the next trial. Participants may control their eye movements in a different manner when they are aware they have a sufficient break in between. One possibility is that their found individual differences are driven mostly by an increasing deficiency to switch back and forth between trial and break – reflecting difficulties in executive functioning (which has been related to ADHD – see Willcutt, Doyle, Nigg, Faraone & Pennington, 2005 for a meta-analysis). To answer this, it would be necessary to investigate individual differences in how oculomotor variability evolves over the time course of individual trials and of the experiment as a whole – rather than looking only at a mean saccadic rate over all trials – to get more insight into possible mechanisms underlying these potential individual differences.

Our results also diverge from Fried et al. (2014), who found increases in microsaccade and blink rates, but not in pupil size variability, in a clinical ADHD population compared to healthy controls. However, again there are profound differences between these two studies: In Fried et al. (2014), participants performed a rapid action selection task with trials of 2 seconds long (Fried et al., 2014), that featured a visual stimulus in each trial – and as such, meant to capture exogenous variability. In this case, the individual differences aim to reflect functional, task-based deficiencies in ADHD patients. As such, their task may be more sensitive to capturing such individual differences.

Overall, our findings show that the benefit of endogenous oculomotor variability as an objective surrogate to self-assessed personality traits in unclear. While we did find some correlations, the correlation coefficients (and Bayes Factors) were small (ranging .22-.26). One may argue that these ranges are typical in Psychology – as Gignac & Szodorai (2016) showed in a meta-analysis of 708 correlation coefficients. However, it is important to note that in our results, the correlations between the different questionnaires were in much higher ranges. Of course, we do not want to deny the importance of finding biomarkers for ADHD. However, within the context of our findings, the benefit of measuring oculomotor activity seems unclear when short and simple questionnaires lead to much larger effect sizes.

One important point to bring up is the severity of symptoms. Our experiments were conducted on healthy participants, and as a result of that, there were not many individuals at the high end of the spectrum. If any individual differences exist, they will be more pronounced when comparing extremer cases. In healthy and academic samples, these more extreme cases will be difficult to find by chance. Comparing clinical cases to healthy controls (such as Fried et al., 2014) will by default be more sensitive to individual differences. In other words, even if our oculomotor measures do not appear beneficial for differentiating between healthy individuals, they may still prove useful to distinguish clinical cases of ADHD or assess the possible benefits of medication.

### Mechanisms underlying potential individual differences

Within the context of our study, we have discussed possible associations between oculomotor variability and ADHD. This may imply that oculomotor variability is inherently detrimental. Of course, this would be a false assumption; oculomotor variability inherently reflects the functioning of our oculomotor system. Fixational eye movements have been proven to be important for our vision (see Rolfs, 2009; Martinez-Conde et al., 2013 for reviews).

Within the context of oculomotor resting states, when participants are instructed to keep fixation, higher variability may be perceived as ‘worse performance’. This means that on the one hand, as ADHD symptomalogy has been associated with decreased task performance in other types of tasks (see Kofler et al., 2013 for a meta-analysis; see Tamm et al., 2012 for a review), it could also be associated with decreased ‘fixation performance’. On the other hand, because fixational eye movements are a healthy phenomenon during fixation, they may be reduced in clinical conditions. This highlights the importance of indicating which mechanisms would drive potential individual differences in variability. Instead, task-based oculomotor variability, in which certain eye movement patterns may be considered as beneficial or detrimental for the task, may be better suited to study these individual differences.

### Oculomotor measures: extraction and correlations

In the current analysis, we used a cut-off of two degrees in amplitude; only microsaccades below this cut-off were counted for the microsaccade rate (similar to Fried et al., 2014; Panagiotidi et al., 2017). However, despite this cut-off being a traditional standard in the literature, it remains somewhat arbitrary. Saccades and microsaccades may represent a continuum, rather than two opposing categories (Otero-Millan, Troncoso, Macknik, Serrano-Pedraza & Martinez-Conde, 2008; Otero-Millan, Macknik, Langston & Martinez-Conde, 2013). We therefore reran our (micro-) saccades analyses without an amplitude cut-off. This measure may capture more of the total variability that participants exhibited. However, without this cut-off, results remained highly similar, and conclusions did not change.

It should be noted that we also used a cut-off for the extraction of blinks: Blinks were computed as missing samples with a maximum of one second – to differentiate blinks from periods of task disengagement (e.g., a participant falling asleep). Similarly, when rerunning our blink-related analyses without the upper-bound cut-off, our findings did not change.

To extract the microsaccades, we used the binocular detection algorithm of Engbert and Kliegl (2003). One feature of this algorithm is that the threshold for detecting a microsaccade is computed for each trial, to adjust for differing amounts of noise between different trials. However, our tasks do not contain any traditional trials, but continuous measurements of 1-4 minutes. This may affect the computation detection threshold due to untypical variability within the ‘trial’, resulting in too lenient thresholds. Still, our microsaccade rate is well in line with previously reported rates using shorter trials. Furthermore, we also used the measures of gaze variability, which may capture the microsaccades as well at the other types of fixational eye movements thus reflecting an overall capacity to fixate.

Previous research has also looked at the associations between task-based oculomotor measures, and found that the six measures (saccade amplitude, microsaccade rate and amplitude, and fixation rate, duration, and size) that they used could be all be captured by one single factor in a Factor Analysis (Poynter et al., 2013) – they interpret this factor as *“Individuals’ eye-movement behavior profiles”*. In our data, this was not the case. Seven out of ten pairs of measures showed evidence against correlation, with support only for some low correlations of pupil size variability with microsaccade and blink rate (r-values of .31 and .24 respectively). The only exception of gaze variability and the horizontal and the vertical dimension, which unsurprisingly are highly similar (r = .82), as they are intended to measure the same construct. Overall, our measures thus shared little to no variance and cannot be captured by one underlying construct. The differences in analysed measures may explain why Poynter et al. (2013) found one underlying construct in their measures, while we did not. Poynter et al. (2013) used three measures related to fixation, and three measures related to (micro-) saccades. Our measures are quite different from Poynter et al. (2013), with only microsaccade rate overlapping, and seem more divergent from each other.

## Conclusion

In the current study, we found that oculomotor variability shows good correlation within individuals both over time and over different conditions. Particularly microsaccade rate, blink rate, and variability of pupil diameter show good reliability – meaning that these measures have the potential to be used as biomarkers. Of course, this begs the question of *what for* they can be used as biomarkers. Our results showed that the between-subject correlations to self-assessed ADHD, mind wandering, and impulsivity were all either absent or very small. In contrast, the questionnaires themselves correlated well with each. Considering the low costs and ease of questionnaires compared to oculomotor data, the benefit of the latter in differentiating between personality traits remains unclear. Still, it is possible that oculomotor measures may serve a function complementing questionnaires. Future research should focus on linking the resting-state oculomotor measures to task-related deficiencies in ADHD or differences in brain structure or integrity, as in these cases, oculomotor measures may serve as an easy and cheap substitute.

## Acknowledgments

We would like to thank Rachel Draper, Laura Daniells, Laura Fleetwood, and Joel Bentley, who collected the data for Experiment 1, as well as Lingsi Zhou, who helped collect the data for Experiment 2, and Natasha Harris, Humairaa Uddin, and Sarah Lethbridge, who helped collecting the data for Experiment 3. We are grateful to Sam Hutton, for sharing his code and giving advice on microsaccade detection.

## Footnotes

1 Note that BF01 (null over alternative hypothesis) can be derived from BF10 (alternative over null) by taking its inverse.

